# BGEN: a binary file format for imputed genotype and haplotype data

**DOI:** 10.1101/308296

**Authors:** Gavin Band, Jonathan Marchini

**Affiliations:** Wellcome Centre for Human Genetics, University of Oxford, Roosevelt Drive, Oxford, OX3 7BN; Department of Statistics, University of Oxford, 24-29 St Giles’, Oxford OX1 3LB

## Abstract

The impact of modern technology on genetic epidemiology has been significant, with studies comprising millions of individuals assessed at tens of millions of genetic variants now becoming common. Studies on this scale provide logistical and analytic challenges starting with the issue of efficiently storing, transmitting, and accessing underlying data. Here we present a binary file format (the BGEN format) that can store both directly-typed and statistically imputed genotype data, and achieves substantial space savings by data compression and the use of an efficient representation for probabilities. We investigate the properties of this format using imputed data from the UK BiLEVE study, demonstrating both storage efficiency, and fast data loading performance on the order of hundreds of millions of imputed genotypes per second. To make using BGEN as easy as possible, we provide a detailed specification and a freely available reference implementation, and we leverage this by developing additional tools including an indexing tool (bgenix) and an R package (rbgen) that permits loading of BGEN-encoded data into the R statistical programming environment. The UK Biobank is one of a number of projects that have used BGEN for release of imputed data, and we expect the format to continue to be widely implemented and used.

## 1 INTRODUCTION

The need to discover and dissect genetic associations at ever finer detail, and advances in the throughput of genotyping and sequencing technologies has driven rapid increases in the scale of genetic epidemiology studies. Studies incorporating millions of individuals with genotypes either directly observed or inferred at tens of millions of genetic variants are now becoming common^1-3^. Data of this scale has effectively outgrown the practical limits of first-generation GWAS data formats^4,5^ and new tools for storage, computation, and analysis are required. For example, the UK Biobank has released data for almost 500,000 individuals with genotypes at over 80 million genetic variants^1^, requiring storing on the order of 1×10^13^ genotypes. Further, many of these genotypes are imputed (probabilistically inferred from directly-typed genotypes using population reference panels^6,7^), necessitating formats that handle imputation uncertainty. Simply storing and accessing these data is thus a major challenge.

Recent work has focused on methods to exploit the linkage disequilibrium structure of the human genome to achieve extremely high compression rates with efficient access to genotypes^8^. These approaches are highly promising but do not currently extend to imputed genotypes.

Here, we focus on the problem of storing large imputed datasets in an efficient way. We present a binary file format (the BGEN format) that is designed to meet specific goals. First, the format should be widely applicable - in particular it should be capable of storing both directly observed and imputed genotype data, should support both unphased genotype and phased haplotype data, and should handle common but sometimes awkward cases such as multiallelic variants and haploid genotype calls. Second, the format should achieve a good balance between storage size and file access performance. Data of this type is often created once but read many times in subsequent analyses and therefore we focus particularly on the efficiency with which data can be read. Third, since a common use case is to access data on specific variants or regions of the genome, the format should be amenable to indexing, effectively permitting random access.

Most importantly we aim for the format to be practically useful - implying in particular that it is widely implementable. To aid with this, we provide a detailed specification of the format (available at http://www.bgenformat.org), a set of supporting tools, and an open-source implementation that is suited for incorporation into other software. Support for BGEN is now available in a number of packages, including QCTOOL, SNPTEST, PLINK^9^, BOLT-LMM^10^, REGSCAN^11^, BGENIE^1^, LDSTORE^12^, MEGA2^13^, and F1AIL^14^ (see URLs). Several major projects including the UK Biobank^1^, ALSPAC^15^, The Human Connectome Project and the MalariaGEN project^16^ have used the BGEN format for data release. Widespread software support for BGEN is thus shortening the path from this data to the insights it can provide.

## 2 METHODS

### The BGEN format

Conceptually, a BGEN file stores genotype probability data for a specified list of samples (with indices *0,…,N-1*, say) and a specific list of variants (with indices *0,…,L-1*). The data for each sample consists of the probability of each possible genotype call that the sample might have at the variant. Here, the possible genotypes are determined by the variant alleles and by the ploidy of the sample (i.e. the number of alleles of the variant that the genome of the sample carries). The ploidy is considered to be known beforehand and is explicitly stored in the file, i.e. BGEN does not specifically handle situations where the ploidy might be unknown or uncertain.

To represent these data in a practically useful way, a BGEN file is organised in a series of data blocks. We now describe these blocks.

First, a *header block* is present which stores metadata needed to properly process the file, including the number of samples and the number of variants, and fields describing the low-level detail of the format being used. Additionally, this block can contain arbitrary data as specified by the user which may be useful e.g. for data versioning.

Second, a *sample identifier block* may be present; this lists an identifier for each of the N samples contained in the file. This block is optional; if not present samples are treated as anonymous.

The remainder of the file consists of a sequence of *variant data blocks*, one for each of the *L* variants stored in the file. Each variant data block contains identifying information about the corresponding variant (e.g. its identifier, genomic position, and alleles), followed by the genotype probability data for the variant. Genotype probability data are encoded in a packed bit-representation, and are then compressed. We describe the layout of this data in subsequent paragraphs. (A simpler version of this data storage, termed BGEN v1.1, is also supported and is described fully in the specification).

### Storage of ploidy and missingness

ploidy for each sample is stored in a sequence of N consecutive bytes. Ploidies from 0 to 63 are supported. Additionally, we use a single bit from each of these bytes to indicate that genotype data for the corresponding sample is missing. To enable efficient processing, we also store the maximum and minimum ploidy across samples; for example this allows implementations to allocate storage efficiently and/or to easily detect the common case where all samples are diploid.

### Storage of unphased genotype data

For a variant with *K* alleles and a sample of ploidy *P* there are

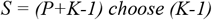

possible unordered genotype configurations. In BGEN these are conceptually enumerated as follows. First, encode each genotype as the tuple of counts of alleles 1, 2, …, K carried by that genotype. (For example, for a biallelic variant and a diploid sample, the possible genotypes are (2,0), (1,1), (0,2) indicating 2, 1, or 0 copies of the first allele). Second, enumerate genotypes in the colex order of these tuples - i.e. in lexicographic order after reading the tuples right-to-left. This definition is compatible with the ordering employed by other common formats, including VCF^17^ and GEN^4^, and applies to arbitrary ploidy and allele counts.

Given a sample of ploidy *P*, let *p*=*(p_1_,…,p_S_)* be the imputed genotype probabilities for the *S* possible genotypes, enumerated in colex order as above. The vector *p* sums to one; in other words it lies on the unit *(S-1)*-simplex Δ_(S-1)_. To reduce the storage requirement for p, we approximate p by another vector in Δ_(S-1)_ that can be encoded using only a specified number of binary bits per probability. Specifically, given a number of bits *b*, we take the closest approximation to *p* on Δ_(S-1)_ that is of the form *q/(2^b^-1)*, where *q*=*(q_1_,…,q_s_)* is a vector with integer coordinates in the range *0, …, 2^b-1^*, i.e. exactly encodable in *b* binary digits. An algorithm for finding *q* was given by Bomze et al^18^. Given *q*, we encode the probability data for the sample by storing *q_1_,…,q_S-1_* in consecutive bits of the file, for a total storage of *b(S-1)* bits. The final value *q_s_* is not stored, since *p_s_* can be inferred as one minus the value of the other probabilities. Probabilities for consecutive samples are encoded in consecutive bits in the file.

The transformation from *p* to *q* introduces rounding error into the stored probabilities. Assuming the probability vector *p* sums to one, we have the error bound |*q-p*|_*1*_ < *1/(2^b^-1)*, i.e. the maximum difference between a stored probability and the true probability is at most 1/(2^b^-1). Thus, the minimum number of bits per probability required for a given level of accuracy *ε* is *b*=*ceil(log_2_(1+1/ε))*. For example, hard-called genotype data can be faithfully stored using *b*=*1*; while the values *b*=*5, 8, 11, 15, 18* give approximately one, two, three, four, and five decimal places of accuracy respectively. (Based on the results below, we recommend using *b*=8 or above for imputed datasets). Unlike schemes that independently round each probability to a specified precision independently, the approximate genotype probabilities stored in BGEN always sum to 1, avoiding potentially subtle numerical issues in downstream analyses.

### Storage of phased genotype (haplotype) data

Phased data is stored as described above, except the approximation takes place for each of the *P* haplotypes carried by the sample. Data for each of the *P* haplotypes is stored consecutively. Full details are provided in the specification. We note that, unlike VCF format, BGEN has no mechanism to identify blocks of phasing; instead, haplotypes are assumed to be in the same order at each variant (i.e. to be globally phased within each chromosome).

### Compression

Finally, all genotype data for the variant, including the ploidy/missingness bytes and the encoded probability values are compressed using either the zlib (http://zlib.net) or zstandard (http://www.zstd.net) compression libraries. The compressed data is stored immediately following the variant identifying data (which is stored uncompressed) and is followed either by next variant data block, or the end of file if no further variants are present.

### Implementation

To make BGEN widely implementable, we developed both a detailed file format reference, and a reference implementation written in C++ (see URLs). This implementation is designed to be easily incorporated into other software; in particular it exposes a flexible interface that avoids imposing specific storage data structures on the user. Several packages developed by other authors have successfully used this library for BGEN support^11-13^.

### Indexing

A design goal of BGEN is to allow indexed access to the compressed data (i.e. to allow efficient access to specific genomic ranges or specific variants). In BGEN, the separation of variant identifying data (stored uncompressed) and compressed genotypes makes this particularly efficient. We illustrate this by implementing a BGEN index file format, based on sqlite3, in the program `bgenix` (URLs). This program behaves similarly to the popular tabix utility for indexing text files^19^, and permits efficient access to genomic ranges or to variants with specific identifiers.

### Interface to R

We illustrate use of the bgen reference library by implementing an R^20^ package (*rbgen*) based on the bgen reference implementation and Repp^21^. This allows the user to directly load probability data from an indexed BGEN file into R without intermediate steps, facilitating a wide range of analyses that are available in R.

### Performance comparison

We assessed the performance of the BGEN format on a dataset of 49,458 samples from the UKBiLEVE study^22^, imputed into the 1000G Phase 3 reference panel. We used data from chromosome 22, which comprised 9,737 directly-typed variants and 521,675 imputed variants. Data was supplied in gzipped GEN format, with probabilities stored to 3 decimal places. We re-encoded the data into BGEN format with 1, 2, 4, 8, 12, 16, and 20 bits per probability, as well as into the plink binary (BED), VCF, and BCF formats. In this comparison, BGEN formats with lower numbers of bits, and BED format lose precision relative to the original data.

We compared the performance of two common, simple operations - reading the variant identifying data without processing genotypes, and reading all the genotype data for each variant into memory without additional processing. For the latter comparison, we used the `–read–test` option of the QCTOOL program to read in all the different data formats. (A caveat is that the QCTOOL implementation of these formats may not be the fastest possible - in particular, plink^9^ has a fast implementation of BED format that is likely to outperform the implementation used here. However, this places all comparisons in a consistent processing framework thus making them comparable.)

Both zlib and zstandard compression methods supported by BGEN also allow tuning via a compression parameter. We compared both the maximum compression available with each library (zlib level 9, zstandard level 21), and an intermediate value chosen to give a reasonable tradeoff between file size and compression speed (zlib level 6, zstandard level 17). All performance comparisons were made using a machine included in the Wellcome Centre for Human Genetics compute cluster, with data stored on the cluster storage filesystem. The test machine had an Intel Xeon CPU clocked at 2.6Ghz, with code compiled using gcc 5.4.0 and avx, sse2, and ssse3 instruction sets turned on. All timing measurements were made using a single thread of execution, and were run in triplicate, with the best of three times reported.

### Assessment of file size scaling with number of samples

To assess scaling of file size with number of samples, we used QCTOOL to subsample the 8-bit encoded data to randomly chosen subsets of between 10 and 31,623 samples.

### Downstream effect of loss of precision

To investigate the potential effect of reduced precision encodings on downstream association analyses, we simulated case/control and quantitative traits and tested for association as follows. First, we simulated five binary traits with different case/control ratios by independently sampling a phenotype for each sample from a Bernoulli distribution, using a success probability of 10%, 25%, 50%, 75%, or 90%. Secondly, we simulated 100 quantitative traits including ‘true’ genetic associations as follows. We grouped variants into five minor allele frequency ranges (1-2%, 2-5%, 5-10%, 10-20%, 20-50%) and three ranges of IMPUTE info score (0.1-0.5, 0.5-0.9, 0.9-1). For each of the 100 traits, we randomly picked an allele frequency and info score bin uniformly among bins, and randomly chose a trait-associated variant from within this bin. We drew the variant effect size from a Gaussian distribution with mean 0 and standard deviation equal to 0.04. We then simulated trait values as *trait = gβ + ε*, where *g* is the vector of genotypes at the associated SNP, and *ε* is chosen to make the overall trait variance equal to 1, i.e. *var(ε)=1-var(gβ)*. We tested for association with case/control traits using SNPTEST and with QTL traits using BGENIE^1^.

We also investigated the effects of the encoding on allele frequencies by computing the expected allele frequency (i.e. the sum of expected genotype dosages divided by twice the number of samples) for each variant under each encoding. We compared the computed expected allele frequency with the ‘true’ expectation, computed using the *b*=16 bit data.

## 3 RESULTS

### File size

BGEN formats with <= 20 bits per probability had lower file sizes than either GEN format (here compressed using zlib level 9, 4.3Gb) or plink binary format (uncompressed, 6.2Gb) (**Fig 1**). While the highest precision BGEN file was comparable in size to the source data (*b*=20 bits per probability, 3.7-4.3Gb), comparable precision in BGEN is achieved using *b*=12 bits, here giving 2.8-3.5Gb, a saving of approximately 60-80% over the source data, depending on compression used. Further decreases in storage size can be achieved at the cost of some loss of precision (e.g. 2.3-2.7Gb at 8 bits per probability, 1.3-1.6Gb at 4 bits per probability). We assess the likely impact of loss of precision on downstream analyses below. The zstandard method of compression had consistently improved file sizes relative to zlib in our comparison (**Fig 1**), despite broadly similar or improved encoding times.

**Fig 1:**
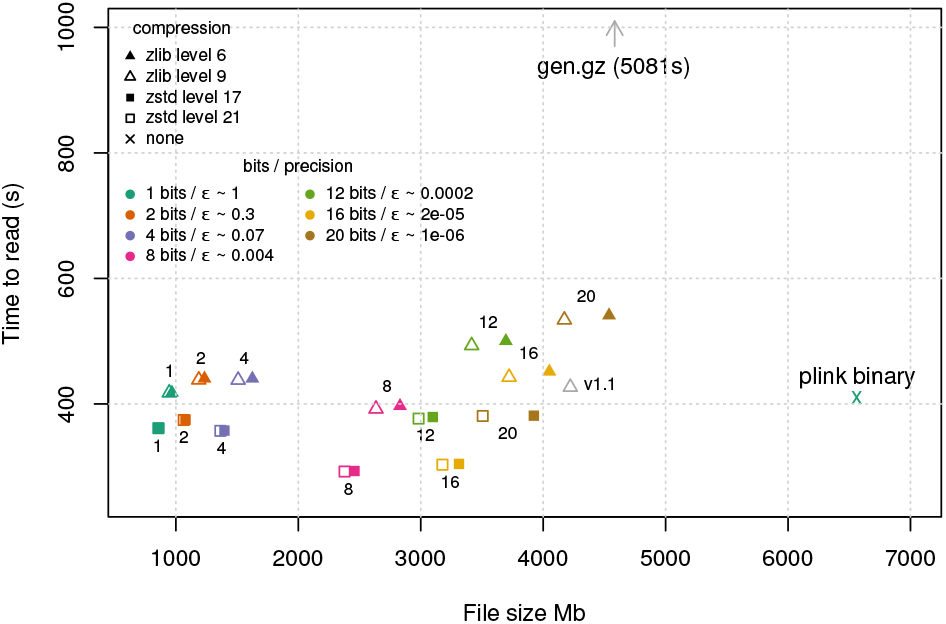
Comparison of file size and data read performance.

### File reading performance

In our file reading speed test, data for each variant in the dataset is parsed and stored in-memory in floating-point form, intended to reflect typical times needed to get data to the point of analysis. Thus overall timings include time for I/O, decompression, parsing, and storage in memory, all of which can be substantial with data volumes of this scale. BGEN formats, as well as plink binary format, achieved much greater performance than the GEN-formatted source data (5,081s), reflecting the fact that binary-encoded genotypes can be parsed much more efficiently than human-readable representations. Decompression time was also noticeable here, with zstandard-compressed files taking around 60-80% of the time to read compared to zlib-compressed files, making the zstandard-compressed BGEN files the fastest in this test. File reading performance was fastest for 8- and 16-bit-encoded data (e.g. 328s for 8-bit encoded data, or ~240 million genotype probabilities parsed per second; 338s for 16-bit encoded data). This is in part because 8- and 16-bit encodings admit particularly efficient parsing implementations; we expect further optimizations to be possible e.g. for multiple-of-4 bit sizes, but do not explore this further here.

The file reading test outlined above involves fully parsing probabilities into floating-point format, and storing in memory. However, another strategy is to attempt computation using the encoded bits directly. To illustrate this, we implemented a simple program to compute the expected allele frequency for each of the 531,412 variants in our test dataset, using the 8-bit-encoded data compressed with zstandard. We implemented this either using the method above, which converts each probability to a floating point number before accumulating, or by using a lookup table to directly map the 16 bits of stored data per-individual to a dosage value. Both programs produced identical output, but the lookup table version was substantially faster, taking approximately 57s to traverse the dataset (i.e. visiting ~460 million genotypes per second) versus around 200s for the floating-point version. Thus, we expect that analysis code that takes advantage of the underlying encoded data will be able to gain additional performance improvements.

### Partioning and scaling of storage size

We observed a roughly linear increase in file size with number of samples (with a limiting slope of approximately 0.05Mb per sample for the zstandard-compressed 8-bit encoded files; **Fig S1**; based on taking subsets of our data). At lower sample counts this trend flattened off, as expected given the constant overhead of storing variant identifying data in each file. We also noted that the lowest confidence and highest-frequency variants occupy more space per variant, on average, than rarer or more confidently imputed variants (**Fig S2**).

### Listing variants and indexing

A particular feature of BGEN is that it stores variant identifying data uncompressed. This choice makes it amenable to indexing by variant and means that accessing variant information without genotypes involves no decompression. In our tests, listing the chromosome, position, ID, and alleles of all 531,412 variants from a BGEN file took approximately 5 seconds. By comparison, the same operation for plink binary format (implemented using the UNIX “`cat`” command applied to the `.bim` file) was essentially instantaneous. Accessing variant identifying data from compressed GEN or VCF files (using zcat and cut), or BCF files (using bcftools^23^) took much longer (>5 minutes), as expected given that this operation involves decompression of the whole file. Our implementation of indexing (bgenix, see URLs), which stores the file location of each variant in an sqlite file, also does not require decompression, and took approximately 75s to build an index in our test.

### Downstream effects of lowered precision

The use of a limited number of bits to store probability leads to rounding errors in approximating probabilities. We investigated the likely effects of this on downstream association analyses by computing association test statistics across all variants in chromosome 22 using each encoding of the data, for five simulated case/control traits and 100 simulated quantitative traits. We compared results for b=1, 2, 4, 8, 12, or 16-bit-encoded data to the full precision data (encoded with b=20 bits; **Fig 2**). Data encoded with b=8 or above showed little difference in association test results (maximum difference in z-score (MDZ) = 0.11, max difference in -log10 P-value (MDP) = 0.09), with smaller differences for more common or better-imputed variants (e.g. MDZ=0.02, MDP=0.03 for variants with info > 0.5 and expected minor allele frequency (MAF) > 0.5%). Similar, but larger discrepancies were observed for b=4 bits (MDZ=1.6, MDP = 1.1 across all variants; MDZ=0.4, MDP = 0.4 for variants with info > 0.5 and MAF > 0.5%).

**Fig 2:**
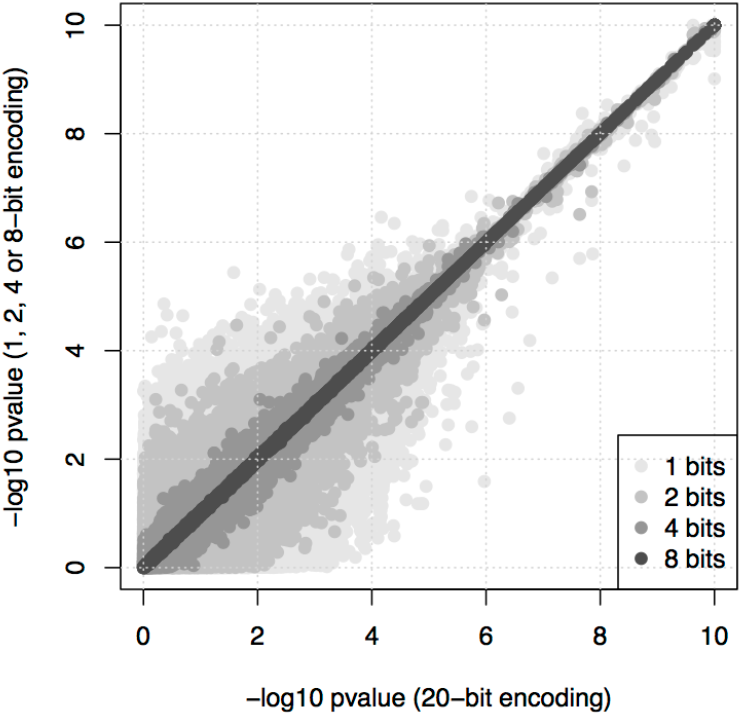
Comparison of association test P-values for simulated case/control and qtl traits

We note that lower-precision encoding has a systematic impact on allele frequency estimates for less confidently imputed variants, an effect that is similar to but less extreme than that caused by thresholding genotype probabilities (**Fig S3**). Specifically, lower-precision data tends to underestimate the expected frequency of rarer alleles. This effect decreases with increasing imputation confidence, with increasing minor allele frequency, and with approximation precision, and is almost not observable for *b*=8 and above.

Given the findings above we recommend the use of b=8, 12 or 16 bits for imputed data, with b=8 providing a particularly good balance between implementation efficiency, accuracy, and overall file size. Directly genotyped or ‘hard called’ data can be losslessly stored using b=1.

## 4 DISCUSSION

The use of ever-larger cohorts typed or imputed at many millions of variants is necessitating new strategies for computation, storage, and data processing. Here we present a binary file format, the BGEN format, that is suitable for storing both typed and imputed genotype data. BGEN has several advantages over traditional text file formats used for this purpose, such as GEN and VCF, including smaller file sizes and substantially more efficient file reading speeds, and supports efficient indexing by variant.

The feature set of BGEN is designed to make it applicable in a variety of situations. However, there are two important ways in which BGEN is limited. First, BGEN can store genotypes and imputed genotype probabilities, but unlike general-purpose formats like VCF or BCF it cannot currently be used to store other values of interest, such as sequence read counts or genotype likelihoods. This may limit the applicability of BGEN in some contexts. Second, BGEN uses the strategy of compressing data for each variant independently. This has the advantage of simplicity of implementation, but methods that can take advantage of sharing of haplotype segments between individuals, such as those based on PBWT^8,24^, effectively model the LD structure of the genome, and as a result are likely to be able to achieve considerably higher compression and, potentially, file access speed. However, these methods are not currently applicable to imputed genotypes.

We have presented some comparative timing data for a typical use case of reading imputed genotype probability data into floating-point variables held in memory. With sample sizes now reaching millions, the high memory and processing requirements of this representation mean it may not be the most efficient way to process data. As we illustrate above, a promising approach is to make direct use of encoded probability values in BGEN, and we expect that future research in this direction could lead to highly efficient analysis code.

Ultimately the measure of a file format may be the extent to which it is used in practice. The design of BGEN presented here was motivated by the needs of the UK Biobank project, which has released imputed genotype data for almost half a million samples in BGEN format^1^. In addition to tools incorporating our reference implementation^1,11-13^, BGEN implementations now exist in other popular genetic analysis software including PLINK^9^, BOLT-LMM^10^, and HAIL^14^. Additionally, we have made a suite of tools available including the ‘rbgen^1^ R package which lowers the barrier to applying the full range of analyses available in R to BGEN-encoded data. Our hope is that this combination of features, implementations, and available data will make this format useful for the next generation of genetic epidemiology studies.

## 5 URLS

The BGEN file format reference is available at: http://www.bgenformat.org

The bgen reference library, bgenix indexing tool, and rbgen R package are available at: http://www.bitbucket.org/gavinband/bgen

QCTOOL is available at: http://www.well.ox.ac.uk/~gav/qctool

## ACKNOWLEDGEMENTS

G.B. is part of the MalariaGEN resource centre, supported by the Wellcome Trust (090770/Z/09/Z; 204911/Z/16/Z). The Wellcome Centre for Human Genetics is supported by the Wellcome Trust (203141/Z/16/Z). J.M. acknowledges funding for this work from the European Research Council (ERC; grant 617306) and the Leverhulme Trust. We thank Colin Freeman and Jerome Kelleher for their help with the software and manuscript.

